# Hypothalamic Sex-Specific Metabolic Shift by Canagliflozin during Aging

**DOI:** 10.1101/2024.03.21.586115

**Authors:** H.S.M. Jayarathne, R. Sullivan, L. Stilgenbauer, L.K. Debarba, A. Kuchumov, L. Koshko, S. Scofield, W. Liu, B.C. Ginsburg, R.A. Miller, M. Sadagurski

**Affiliations:** Department of Biological Sciences, Bio (Integrative Biosciences Center), Wayne State University, Detroit, MI; Department of Pharmaceutical Science, Bio (Integrative Biosciences Center), Wayne State University, Detroit, MI; Institute of Environmental Health Sciences, iBio (Integrative Biosciences Center), Wayne State University, Detroit, MI; Department of Psychiatry and Behavioral Sciences, University of Texas Health Science Center, San Antonio, TX; Department of Pathology and Geriatrics Center, University of Michigan, Ann Arbor, MI

**Keywords:** Canagliflozin, brain, hypothalamus, metabolism, longevity

## Abstract

The hypothalamus undergoes significant changes with aging and plays crucial roles in age-related metabolic alterations. Sodium-glucose co-transporter 2 inhibitors (SGLT2i) are anti-diabetic agents that promote glucose excretion, and promote metabolic homeostasis. Recent studies have shown that SGLT2i, Canagliflozin (Cana), can extend the median survival of genetically heterogeneous UM-HET3 male mice and improve central metabolic control via increases in hypothalamic insulin responsiveness in aged males, as well as reduced age-associated hypothalamic inflammation. We studied the long- and short-term effects of Cana on hypothalamic metabolic control in UM-HET3 mice. We show that Cana treatment reduced body weight and fat mass in male mice that was associated with enhanced glucose tolerance and insulin sensitivity observed by 12 months. Indirect calorimetry showed that short-term Cana treatment (5 months) increased energy expenditure in male, but not female mice, at 12 months of age. Long-term Cana treatment (18 months) increased metabolic rates in both sexes, and markedly increasing formation of both orexigenic and anorexigenic projections to the paraventricular nucleus of the hypothalamus (PVH) mostly in females by 25 months. Hypothalamic RNA-sequencing analysis revealed sex-specific genes and signaling pathways related to insulin signaling, glycogen catabolic pathway, neuropeptide signaling, and mitochondrial function upregulated by Cana, with males showing a more pronounced and sustained effect on metabolic pathways at both age groups. Overall, our data provide critical evidence for sex-specific mechanisms that are impacted by Cana during aging suggesting key targets of hypothalamic Cana-induced neuroprotection for metabolic control.

## Introduction

The aging process is accompanied by various physiological changes, including alterations in hypothalamic function. The hypothalamus, a crucial brain region that serves as a master regulator of energy expenditure, body temperature, sleep-wake cycles, hunger and satiety, thirst, and hormone secretion (Williams et al., 2001). Age-related changes in the hypothalamus can lead to a cascade of physiological dysfunctions that contribute to the onset and progression of metabolic diseases (Coll and Yeo, 2013).

Emerging evidence suggests that anti-aging pharmacological interventions have the potential to positively impact whole-body metabolic health, thereby exerting beneficial effects on the hypothalamus during the aging process (Liu et al., 2022). One notable class of drugs is sodium-glucose cotransporter 2 inhibitors (SGLT2i) (Xu et al., 2021). These medications were initially developed to manage T2D by reducing blood glucose levels independent of insulin, but recent studies highlight that their beneficial effects extend beyond glycemic control (Wong et al., 2021). We have recently shown that SGLT2i, Canagliflozin (Cana), extends the median survival of male mice by 14%, and robustly retards age-related lesions in male mice without an effect on females (Miller et al., 2020; Snyder et al., 2022). Moreover, long-term Cana treatment led to a significantly lower fasting blood glucose levels and improved glucose tolerance by 22 months of age in both males and females (Miller et al., 2020). SGLT2i cross the blood-brain barrier (BBB) and co-transporters are present in various parts of the brain (Koepsell, 2020; Tahara et al., 2016). Indeed, treatment with the SGLT2i empagliflozin, for 8 weeks, of pre-diabetic patients restored hypothalamic insulin sensitivity, showing the beneficial effects of SGLT2i (Kullmann et al., 2021). The beneficial effects of SGLT2i on overall metabolic homeostasis appear to rely on intact brain-periphery cross-talk via the parasympathetic nervous system (Sawada et al., 2017).

We have previously demonstrated that Cana impact on the hypothalamus of aged mice is sex-specific. For example, Cana treatment significantly improved insulin responsiveness in the hypothalamus of aged male but not female mice. Furthermore, Cana treatment improved exploratory and locomotor activity of 30-month-old male but not female mice. However, both Cana-treated male and female mice showed significant reductions in age-associated hypothalamic inflammation (Jayarathne et al., 2022). The molecular mechanism underlying the sex-specific neuroprotective effects of Cana on hypothalamic regulation of metabolism during aging are unclear.

In this study, we dissected the impact of Cana treatment on hypothalamic transcriptional remodeling during aging, using the genetically heterogeneous UM-HET3 mice to avoid the one-genome effect and replicate to some degree the variability seen in the human population (Bou Sleiman et al., 2022).

## Materials and Methods

### Mouse husbandry

Mice were bred as the progeny of (BALB/cByJ x C57BL/6J)F1 mothers (JAX #100009) and (C3H/HeJ x DBA/2J)F1 fathers (JAX #100004), so that each mouse is genetically unique and a full sibling to all other mice with respect to segregating nuclear alleles (Jackson et al., 1999). Mice are housed at three males or four females per cage from weaning and are provided food (Purina 5LG6) and water ad libitum. Cana was administered in the feed at 180 ppm beginning at 7 months of age until sacrifice at 12 and 24 months of age as previously described (Miller et al., 2020). All mice were provided with water ad libitum and housed in temperature-controlled rooms (22LC) on a 12-hour/12-hour light-dark cycle. If animals exhibited any indication of illness or distress, the laboratory staff conferred with on-site veterinary staff immediately to recommend appropriate interventions. Anesthesia for euthanasia was by avertin. The health status checks were conducted regularly in the animal breeding facility. All animal experiments were performed in accordance with NIH guidelines for Animal Care and Use, approved and overseen by Wayne State University Institutional Animal Care and Use Committee (IACUC).

### Indirect calorimetry

In order to assess VO_2_ consumption, VCO_2_ production, respiratory exchange ratio (RER) and heat production mice were placed in PhenoMaster metabolic cages (PhenoMaster, TSE system, Germany #160407-03). Animals were individually housed during 12 hours of light and dark cycle (Dark cycle: 6 PM – 6 AM; light cycle: 6 AM-6 PM). The mice were acclimatized for 48h and data was collected for 72h while food and water were provided *ad libitum*.

### Glucose tolerance test

For glucose tolerance test (GTT), mice were fasted for 6 hours and intraperitoneally injected with D-glucose at a dose of 2 g/kg⋅BW. Blood glucose levels were measured at basal state (0 min) and at 15, 30, 60, 90, and 120 minutes after injection. Blood glucose levels were measured at the indicated times via tail vein bleeding using OneTouch glucometer. Blood insulin was determined on serum from tail vein bleeds using Insulin ELISA kit (Crystal Chem. Inc.).

### Perfusion and immunolabeling

Perfusion and immunolabeling are performed as previously described (Lima J.B.M., 2020). Briefly, mice were anesthetized and perfused using phosphate buffer saline (PBS) (pH 7.5) followed by 4% paraformaldehyde. Brains were post-fixed, dehydrated, and then sectioned coronally (30 μm) using a sliding microtome, followed by immunofluorescent analysis. For immunohistochemistry brain sections were washed with PBS six times, blocked with 0.3% Triton X-100 and 3% normal donkey serum in PBS for 2 h; then the staining was carried out with the following primary antibodies overnight: rabbit anti-AgRP (1:1000; Phoenix Pharmaceuticals Inc; Cat.No.H-003-53), rabbit anti α-MSH (1:1000; Phoenix Pharmaceuticals Inc; Cat. No. H-043-01), rabbit anti-CART (1:2000; Phoenix Pharmaceuticals Inc.; Cat. No. H-003-62). Brain sections were incubated with Alexa Fluor-conjugated secondary antibodies for 2 h (Invitrogen). Microscopic images of the stained sections were obtained with a Nikon 800 fluorescent microscope using Nikon imaging DS-R12 color cooled SCMOS, version 5.00.

### Hypothalamic RNA extraction

Male mice were sacrificed at *ad libitum* to harvest the brain and to isolate the hypothalamus. Hypothalamus samples were lysed with 0.75 ml of 2-mercaptoethanol added lysis buffer (PureLink® RNA Mini Kit #12183025). Following the homogenization, with 0.75 ml of 70% ethanol, samples was transferred to spin cartridges with the collection tubes. Samples were centrifuged at 12000 x g for 15 sec at room temperature. After washing the samples three times with washing buffers, cartridges were centrifuged at 12000 x g for 1-2 min to dry the membrane with bound RNA. RNA was eluted using 15-20 ul of RNase-free water.

### RNA sequencing and data analysis

Hypothalamic RNA-Seq was performed at the WSU Genome Sciences Core. RNA concentration was determined by NanoDrop (ND-1000 UV-Vis Spectrophotometer) and quality was assessed using RNA ScreenTape on a 4200 TapeStation. RNA-seq libraries were prepared according to the QIAseq Stranded RNA Library Kits protocol before sequencing on a NovaSeq 6000 (2 x 50 bp). Hypothalamic RNA sequences were mapped to the mouse reference genome (GRCm38.90) using HISAT2 v.2.1.0.13 following the adapter trimming and quality checking. Quantification of the gene expression was generated using HTSeq-counts v0.6.0. The male and females were analyzed separately and following a PCA analysis of the raw data clear outliers were removed. Significantly differentially expressed genes (DEGs) were generated using the R Studio package DESeq2. Statistical significance was calculated by adjusting the P values with the Benjamini-Hochberg’s false discovery rate (FDR). All the noncoding RNA was removed from the analysis. DEGs were used to identify the enriched pathways, both Gene Ontology (for Biological Processes (BP)) and KEGG enrichment pathways using Gene Set Enrichment Analysis (GSEA) (NIH DAVID Bioinformatics https://david.ncifcrf.gov/home.jsp) (p-value cut off <0.05). Heatmaps, bubble plots, Venn diagrams and gene expression changes were plotted using SRplot (https://bioinformatics.com.cn/). All RNA-Seq data are available at the Sequence Read Archive (SRA) at NCBI under accession number PRJNA1089563

### Statistical analysis

Results are expressed as the mean ± standard error and were analyzed using Statistica software (version 13.5.0.17). Graphs were generated using GraphPad Prism software. Sample size is annotated within figure legends. All parameters were analyzed by two-way ANOVA using the general linear model function and a full factorial model, which included an effect of treatment, the effect of sex, and the interaction effect between sex and treatment followed by Newman–Keuls post hoc test. The level of significance (α) was set at 5%.

## Results

### Age- and sex-dependent changes in energy homeostasis in response to Cana feeding

We subjected 7-month-old genetically heterogeneous UM-HET3 mice to a Canagliflozin (Cana)-containing diet, as previously detailed (Miller et al., 2020). After 4-weeks of treatment, Cana-fed male mice displayed a noticeable reduction in body weight and fat mass, while Cana-fed females did not show changes in body weight or body composition (Supplementary Fig. 1A-B). In our previous study we found that by the age of 12 months, Cana feeding led to a significant decrease in body weight in both sexes (Miller et al., 2020). Both Cana-fed males and females exhibited slightly enhanced glucose tolerance, but only Cana-fed males showed increased insulin sensitivity, as evidenced by HOMA-IR (Supplementary Fig. 1C-F). To further assess the effect of Cana on metabolic control during aging, we measured parameters of whole-body energy homeostasis by indirect calorimetry. Oxygen consumption (VO_2_) and carbon dioxide production (VCO_2_) were significantly elevated throughout the dark and light cycle (p<0.05 for effect of drug) only in Cana-fed 12-month-old males compared to females with a significant interaction between sex and drug treatment compared to control (p<0.001 for light cycle, p<0.05 for dark cycle) (Fig. 1A-L). Moreover, 12-month-old Cana-fed males exhibited a significant increase in energy expenditure (EE) during both light and dark cycles (p<0.05 for effect of drug), without an effect on females with a significant interaction between sex and drug treatment compared to control (p<0.01 for light cycle, p<0.05 for dark cycle) (Fig. 1M-R). There were no significant differences in respiratory exchange ratio (RER) and activity levels between controls or Cana-treated mice (Supplementary Fig. 2A-L). However, by 25 months of age, both aged Cana-fed male and female mice exhibited higher metabolic rate with increased EE and RER, particularly during the dark cycle (p<0.001 for effect of drug) (Fig. 2 and Supplementary Fig. 3A-L). Cana-fed females demonstrated higher RER over the 24-hour period (p<0.01), indicative of an enhanced reliance on carbohydrates as fuel utilization. These changes in RER and energy expenditure were not associated with alterations in activity levels in 25-month-old Cana-fed mice (Supplementary Fig. 3M-R). 12- and 25-months-old Cana-fed mice exhibited significantly increased food consumption and water intake (p<0.001 for effect of drug) (Supplementary Fig. 2M-P and 3S-V). Thus, prolonged Cana feeding supported improvement in energy homeostasis in aged mice of both sexes.

**Figure 1:**
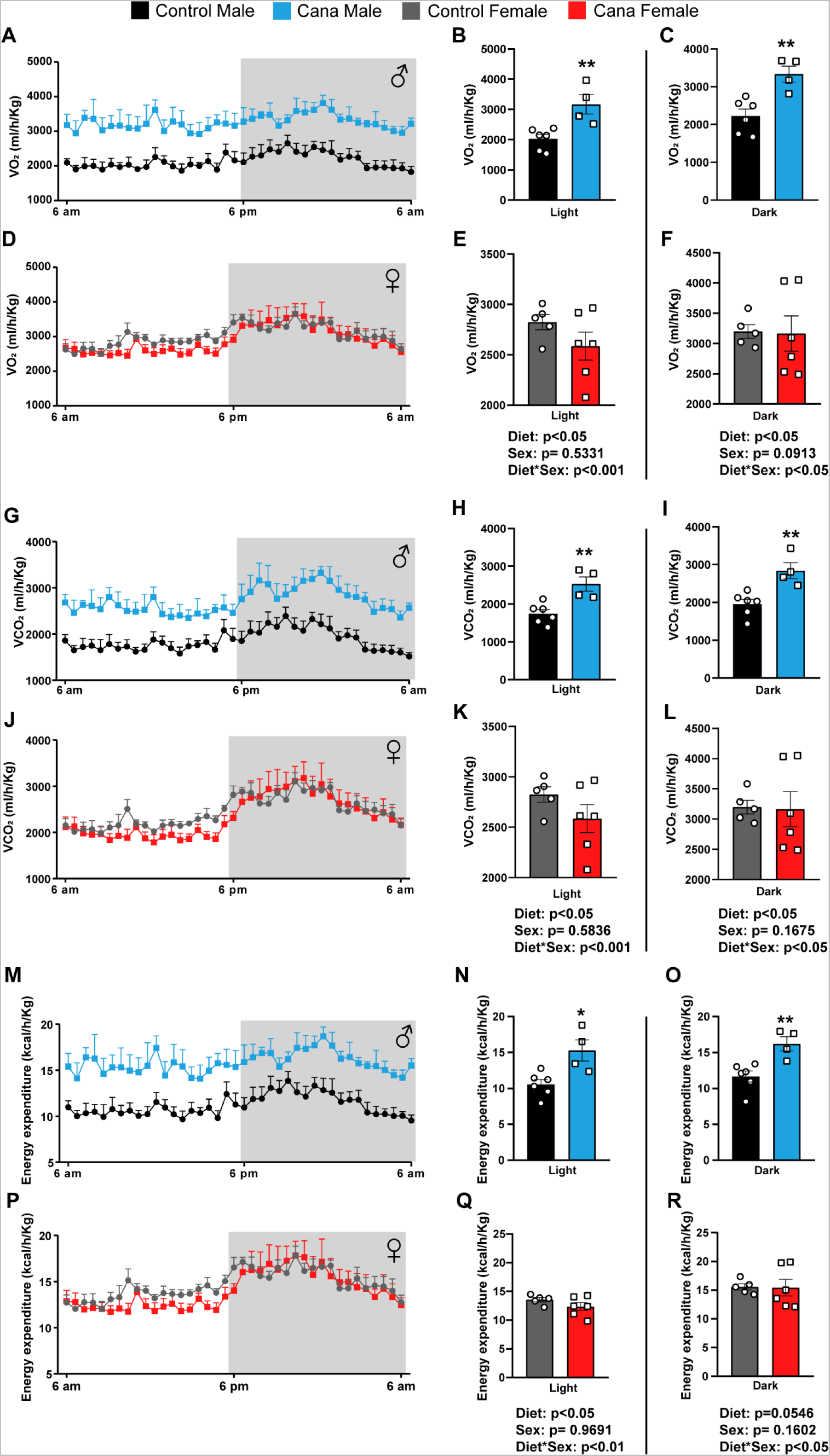
Cana feeding alters energy homeostasis parameters in 12-month-old male but not in female mice. (A-F) Oxygen consumption (VO_2_) in males (A-C) and females (D-F). (G-L) Carbon dioxide production (VCO_2_) in males (G-I) and females (J-L). (M-R) Energy expenditure in males (M-O) and females (P-R). Data represented as mean ± SEM, n= 5-6 mice/group. Two-way ANOVA followed by Newman-Keuls analysis (B-C, E-F, H-I, K-L, N-O and Q-R). *p<0.05, **p<0.01. P values for the effect of diet, sex and the interaction during light or dark cycle represent the significant p values from the two-way ANOVA.

**Figure 2:**
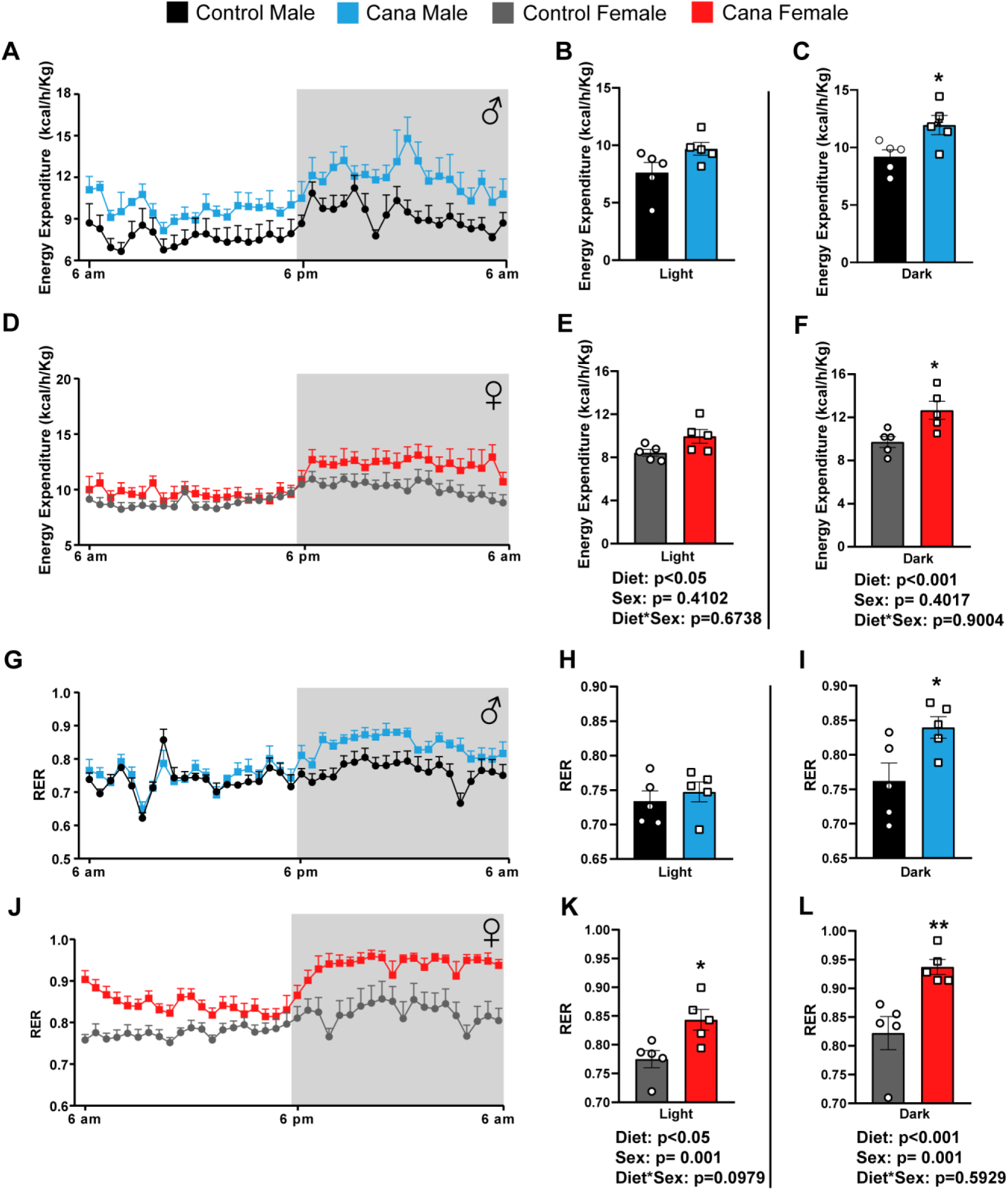
Cana feeding modulates energy homeostasis parameters in aged 25-month-old mice of both sexes. (A-F) Energy expenditure in males (A-C) and females (D-F). Respiratory exchange ratio (RER) in males (G-I) and females (J-L). Data represented as mean ± SEM, n= 5-6 mice/group. Two-way ANOVA followed by Newman-Keuls analysis (B-C, E-F, H-I and K-L). *p<0.05, **p<0.01. P values for the effect of diet, sex and the interaction during light or dark cycle represent the significant p values from the two-way ANOVA.

### Cana feeding alters hypothalamic orexigenic and anorexigenic projections in aged mice

Cana levels in the hypothalamus and cortex were measured using high-performance liquid chromatography/mass spectrometry as before (Jayarathne et al., 2022). Interestingly, Cana levels in the hypothalamus of the females were significantly higher than males (0.36-1.25 ng/mg for females versus 0.12-0.38 ng/mg for males) (Fig 3A), while Cana levels in the cortex were similar between males and females (Fig 3B). To examine the effect of Cana on hypothalamic neuropeptides following treatment, we analyzed the immunoreactivity of agouti-related peptide (AgRP), alpha-melanocyte stimulating hormone (α-MSH) a product of the proopiomelanocortin (POMC) and cocaine- and amphetamine-regulated peptide (CART) containing fibers in the paraventricular nucleus (PVH) of the hypothalamus in 25-month-old mice. The density of orexigenic AgRP-IR fibers notably increased in the PVH of Cana-treated females, but not males, as compared to controls (p<0.05 for effect of drug, p=0.001 for effect of sex) (Fig. 3C and D). Additionally, the fiber density of anorexigenic α-MSH-IR was higher in Cana-treated females, but not males (p<0.05 for effect of drug), with a significant interaction between sex and drug treatment (p<0.001) (Fig. 3E and F). Interestingly, we observed significant upregulation in CART-IR in both Cana-treated males and females as compared to control mice (p<0.001 for effect of drug) (Fig. 3G and H).

**Figure 3.**
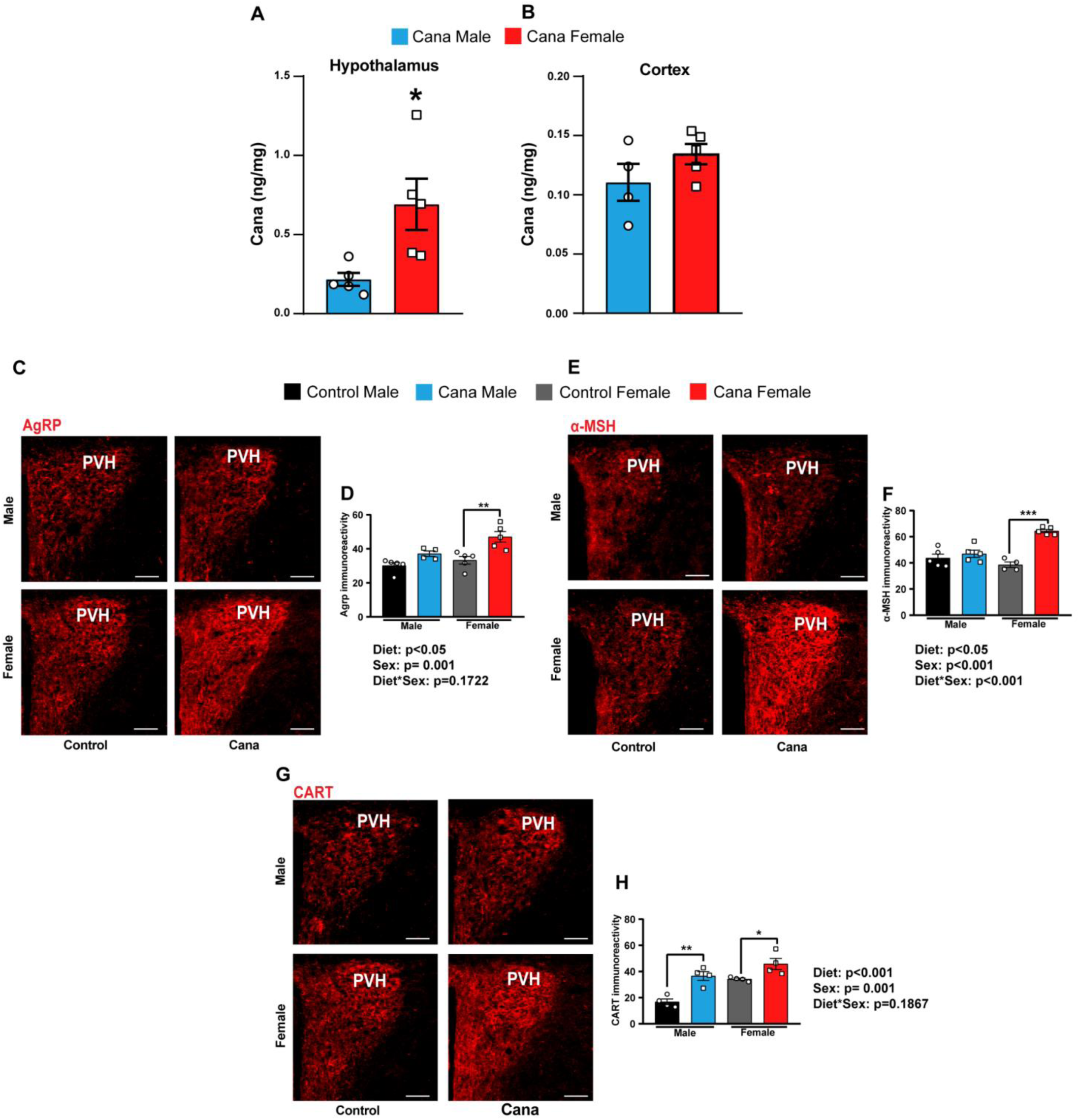
Cana feeding alters hypothalamic projections in aged 25-month-old mice. Cana levels measured in the hypothalamus (A) and cortex (B) in males (blue) and females (red), n=4-5 mice/group. Images and quantification of (C-D) AgRP, (E-F) α-MSH and (G-H) CART immunoreactive fibers innervating the paraventricular nucleus of the hypothalamus (PVH) at 25 months of age in control and Cana treated male and female mice. Scale bar: 200 µm. Data represented as mean ± SEM, n= 4-5 mice/group. Two-way ANOVA followed by Newman-Keuls analysis (D, F and H). *p<0.05, **p<0.01, ***p<0.001. P values for the effect of diet, sex and the diet x sex interaction represent the significant p values from the two-way ANOVA.

### Cana feeding modulates the hypothalamic transcriptome

To gain further insight on hypothalamic responses to Cana during aging, we performed bulk RNA-seq of microdissected hypothalami from Cana-treated males and females at 12- and 25-months of age (Fig. 4). The principal component analysis (PCA) plots demonstrated a clear separation in the gene expression profiles between Cana-treated male and female mice in both age groups (Supplementary Fig. 4).

**Figure 4.**
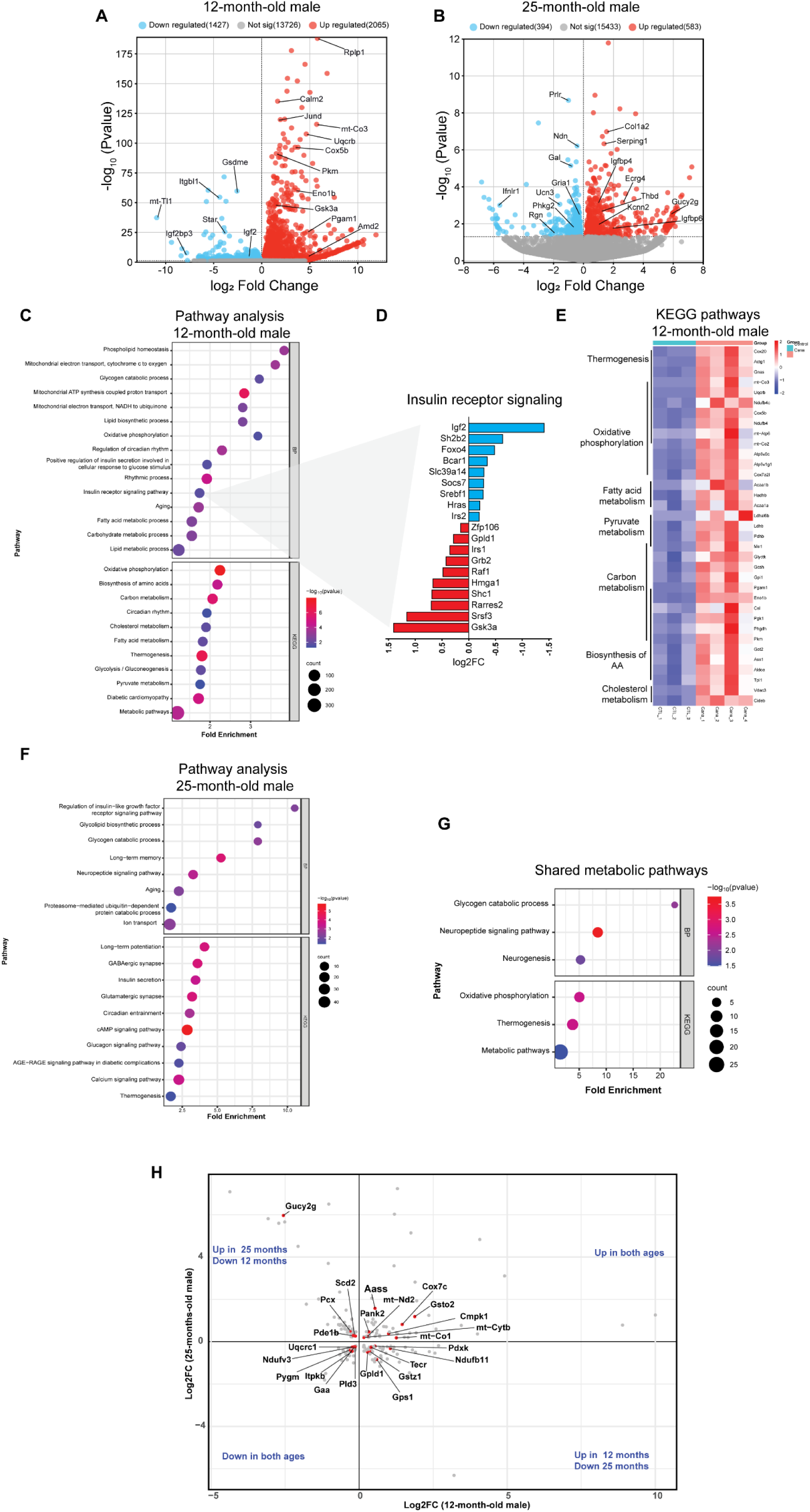
Modulation of hypothalamic transcriptome by Cana in male mice at 12 and 25- months. Volcano plots display differentially expressed genes (DEG, p<0.05), with upregulated genes in red and downregulated genes in blue. (A) Represents 12-month-old, and (B) 25- month-old Cana versus control male mice. (C) Gene ontology (GO) analysis for biological processes (BP) and KEGG pathway analysis in 12-month-old Cana males. (D) Genes related to the BP “Insulin receptor signaling” significantly up and downregulated in 12-month-old Cana males. (E) Heatmap of top upregulated genes in KEGG pathways for 12-month-old Cana males. n = 3-4 mice/group. (F) GO analysis for BP and KEGG pathway analysis for 25-month-old Cana males. (G) Enriched transcriptomic pathways shared between 12 and 25 month-old males. (H) Scatter plot for shared metabolic genes between 12 and 25-month Cana males.

#### Hypothalamic transcriptome in male mice

The volcano plots display genes upregulated and downregulated by Cana in the hypothalamus of male mice in both ages (Fig. 4A and B). We identified 3492 differentially expressed genes (DEGs) in hypothalami of 12-months-old Cana-treated males and 977 DEGs in 25-months-old Cana-treated males, with 196 genes shared between both age groups. Using Gene Set Enrichment Analysis (GSEA), we analyzed the Gene Ontology (GO-BP) and Kyoto Encyclopedia of Genes and Genomes (KEGG) databases to identify biological processes regulated by DEGs. GSEA analysis revealed upregulation of GO-BP terms related to ‘phospholipid homeostasis’, ‘oxidative phosphorylation’, ‘circadian rhythm’, and ‘insulin receptor signaling’ in Cana-treated 12-month-old males (Fig. 4C and Supplementary file 1). Figure 4D illustrates top genes in insulin receptor signaling modified by Cana in 12-month-old males. These genes include downregulated *IGF2*, *Foxo4, Irs2*, and *Sh2b2*, and upregulated *Irs1, Shc1* and *Gsk3a* among the others (Fig. 4D). KEGG analysis showed significant increases in metabolic pathways including carbon metabolism, thermogenesis, oxidative phosphorylation, cholesterol metabolism, fatty acid metabolism, and pyruvate metabolism (Fig. 4E). Examining the impact of hypothalamic aging under Cana, we observed upregulation of metabolic related pathways such as ‘insulin/IGF1 signaling’, ‘glycogen catabolic pathway’, ‘long-term memory’, ‘neuropeptide signaling’ and ‘thermogenesis’ among others in 25-month-old males (Fig. 4F and Supplementary file 1). Notably, GSEA of top overlapping hub genes as showed by scatter plot revealed shared enrichment of metabolic pathways such as ‘glycogen catabolic pathway’, ‘neuropeptide signaling’, ‘oxidative phosphorylation’, and ‘thermogenesis’ in males treated with Cana, for both age groups (Fig. 4G and H). These metabolic pathways commonly modified by Cana suggest their contribution to Cana-mediated hypothalamic control of metabolism in male mice.

#### Hypothalamic transcriptome in female mice

Figures 5A and B illustrate volcano plots displaying the upregulated and downregulated genes induced by Cana in the hypothalamus of female mice for 12 and 25 months of age. We identified altered genes enriched in GO terms in response to Cana treatment in female mice in both age groups. In 12-month-old females (Fig. 5C), these included terms related to metabolic processes such as ’positive regulation of fatty acid activation’ and ’positive regulation of PI3-kinase activity’, as well as pathways associated with ‘aging’, ‘circadian rhythms’, and ‘neurogenesis’. In contrast, 25-month-old females (Fig. 5D) exhibited enrichment in GO terms related to oxidative stress (’positive regulation of reactive oxygen species metabolic process’), as well as signaling pathways such as ’Ras signaling’ and ’PI3K-Akt signaling pathway’. A heatmap illustrates the top genes in the PI3K-Akt signaling pathway affected by Cana treatment in 25-month-old females (Fig. 5E). These genes including upregulated *Fgf22, Creb5, IL7, Tnc, Ngfr and Pdgfb* possess broad cell survival activities. Supplementary file 2 shows the complete GO term and KEGG terms for females. When clustering the top genes with shared enrichment that were affected by Cana in females for both age groups, the clustered genes were primarily related to cellular function and development (Fig. 5F and G). Notably, no overlapping genes related to metabolic pathways were identified, indicating distinct age-related changes in female’s hypothalamic transcriptome affected by Cana.

**Figure 5.**
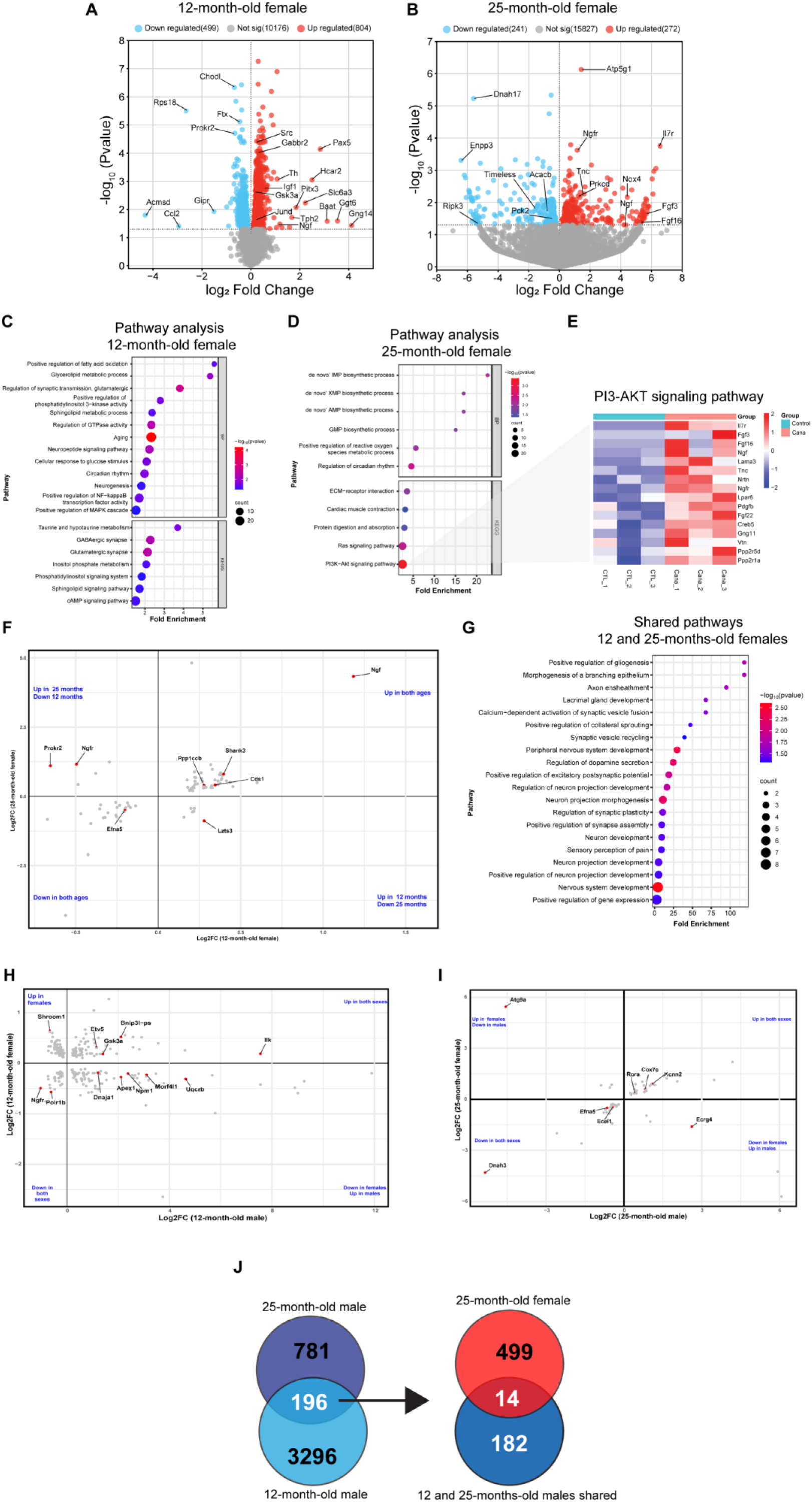
Modulation of hypothalamic transcriptome by Cana in male and female mice at 12 and 25-months. Volcano plots display differentially expressed genes (DEG, p<0.05), with upregulated genes in red and downregulated genes in blue. (A) Represents 12-month-old, and (B) 25-month-old Cana versus control female mice. Gene ontology (GO) analysis for biological processes (BP) and KEGG pathway analysis in (C) 12-month-old and (D) 25-month-old Cana females. (E) Heatmap of top upregulated genes in “PI3-Akt signaling pathway” for 25-month-old Cana females. (F) Scatter plot for shared genes between 12 and 25-month Cana females. (G) Enriched transcriptomic pathways shared between 12 and 25 month-old females. n = 3-5 mice/group. Scatter plot for shared genes between (H) 12-month-old and (I) 25-month-old Cana males and females. (J) Venn diagram depicting the shared genes between Cana-treated males and 25-month-old females.

#### Effect of sex in response to Cana

Finally, we explored the interaction between genes that were differentially expressed by sex in response to Cana. When comparing the transcriptomic changes of the hypothalamus upon Cana treatment in 12-month-old males and females, we observed no shared pathways between males and females, with only a few genes in these pathways exhibiting similar up- or downregulation. For example, the strongest upregulated shared genes *Gsk3a* and *Bnip3l-ps* were related to the ‘insulin signaling’ and ‘mitophagy’ pathways (Fig. 5H). Comparing the transcriptomic changes of the hypothalamus in 25-month-old males and females, only a few genes in shared pathways including ‘thermogenesis’ (*Cox7c*)’ and ‘neuropeptide signaling’ (*Ecrg4* and *Ecel1*) were similarly up- or downregulated (Fig. 5I). Moreover, among the limited number of genes (14 genes) shared among males in both age groups and 25-month-old females, a subset were identified to be related to mitochondrial function and metabolic regulation, such as *Cox7c, Hbb-bs, Gal3st3, Oip5os1,* and *Golga4*, suggesting their specific role in response to Cana (Fig. 5J). Supplementary file 3 shows the complete DEG set for both ages and sex.

## Discussion

The present study highlights the age- and sex-dependent effects of Cana treatment on hypothalamic control of metabolism during aging, with males demonstrating significant metabolic improvements early on compared to females. Additionally, transcriptomic analysis delineates distinct Cana-responsive pathways in each sex, indicating different regulatory mechanisms governing hypothalamic metabolic homeostasis in response to Cana during aging.

Part of the anti-aging effects of SGLT2is relates to their neuroprotective effects (Hierro-Bujalance et al., 2020; Jayarathne et al., 2022; Lin et al., 2014; Naznin et al., 2017). In support, our current results highlight the effectiveness of Cana in the hypothalamus in mitigating age-related metabolic dysregulation in male mice. The early-onset and sustained improvements in metabolic parameters, including reduced body weight, fat mass, and enhanced glucose tolerance in aged UM-HET3 mice of both sexes, are consistent with the known mechanisms of action of Cana in improving peripheral insulin sensitivity and glucose metabolism (Sawada et al., 2017; Xu et al., 2021). The significant alterations in hypothalamic gene expression observed in Cana-treated males in both age groups, particularly related to insulin signaling and neuropeptides signaling, align with our observations that Cana significantly improved central insulin sensitivity in the hypothalamus of aged male but not female mice (Kullmann et al., 2021). Since hypothalamic insulin signaling plays a critical role in regulating energy homeostasis through modulation in neuropeptides signaling pathways (Evans et al., 2014; Spanswick et al., 2000; Xu et al., 2005), this may explain the robust impact of Cana on energy expenditure in male mice. On the other hand, female mice exhibited a delayed response to Cana treatment, with energy homeostasis improvements becoming evident only by 25 months of age. This delayed response in females may reflect sex-specific mechanistic differences observed in Cana mice as well as differences in Cana sensitivity, highlighting the need for further investigation into the underlying mechanisms driving these sex-dependent effects.

In aged mice, there is a significant reduction in AgRP and POMC neuropeptide neuronal activity (Yang et al., 2012), while the age-related metabolic dysfunction can be ameliorated by restoring the POMC gene in the arcuate nucleus (ARC) (Li et al., 2005). Although no genes related to metabolic pathways were identified in 25-month-old females, Cana treatment notably increased the density of AgRP and α-MSH-containing fibers exclusively in female mice compared to controls. Conversely, both Cana-treated male and female mice exhibited higher densities of anorexigenic CART-containing fibers in the PVH, accompanied by increased food and water intake. This supports the concept that SGLT2is induce weight loss partly through urinary calorie excretion, while simultaneously influencing central reward and satiety circuits, leading to increased appetite and food intake (van Ruiten et al., 2022). While both CART and α-MSH neurons are involved in regulating food intake and energy balance, they exhibit distinct patterns of activation and regulation (Lau and Herzog, 2014). These neurons may further exhibit distinct sensitivities to the pharmacological effects of Cana, leading to differential alterations in their responsiveness, suggesting complex sex-specific responses to Cana treatment in the regulation of appetite and energy balance within the hypothalamus.

Sexual differences in responses to Cana might be attributed to differences in Cana pharmacodynamics because we detected a significant effect of sex in Cana levels, with females having higher Cana levels in the hypothalamus, and in the hippocampus than males (Jayarathne et al., 2022). These data are consistent with previous report showing higher levels of Cana in the whole brain of females compared to males (Miller et al., 2020). The observed sex-specific effects of Cana in the hypothalamus might suggest that Cana produces benefits through interaction with a metabolic or physiological pathway particularly relevant to male hypothalamus, or that Cana produced some side effects within the hypothalamus in females. Interestingly, a better response to SGLT2i was reported in men than in women (Dave et al., 2019; Han et al., 2018). Further, sex-differences in *Sglt2* were previously observed in various tissues with males expressing significantly higher levels of *Sglt2* in the whole brain, compared to females (Nagai et al., 2014). The drug effect can be attributed to expression of *Sglt2* in different cell types and regions. Further studies are needed to determine Cana pharmacological effects on the hypothalamic function of both sexes.

Our comprehensive analysis of the hypothalamic transcriptome in response to Cana treatment reveals sex-dependent molecular alterations underlying Cana metabolic impact in aging. Cana treatment elicits distinct gene expression profiles in male and female mice, with notable differences observed between age groups. In male mice, Cana induces significant upregulation of metabolic pathways, including phospholipid homeostasis, oxidative phosphorylation, and insulin receptor signaling, at both 12 and 25 months of age. These findings suggest a sustained metabolic response to Cana treatment and underscores the age-independent effects of Cana on hypothalamic gene expression in male mice. Conversely, female mice exhibit a less pronounced response to Cana treatment, with fewer differentially expressed genes identified compared to males. While Cana induces alterations in pathways related to fatty acid metabolism, PI3K-Akt signaling, and neuropeptide signaling in females, the overall response was weaker and less consistent in gene clusters across age groups. The comparison of conserved genes among males at both age groups and aged 25-month-old females, despite exhibiting phenotypically similar metabolic effects on energy homeostasis, revealed only a limited set of genes with roles in various aspects of cellular metabolism (Grundy, 2004; Iorio et al., 2015; Short et al., 2023; Siklar et al., 2023; van Gerwen et al., 2023; Zhuang et al., 2023). These findings suggest that while aged males and females may exhibit similar metabolic responses to Cana treatment, the underlying central regulatory mechanisms and pathways driving these responses differ.

Overall, our study provides insights into the age- and sex-dependent effects of Cana treatment on hypothalamic control of metabolism. By revealing distinct sex-specific hypothalamic transcriptomic profiles in aged mice, we demonstrate the complexity of Cana’s impact on energy homeostasis regulation in aging. Understanding the central sex-specific mechanisms behind these differences will not only enhance our understanding of metabolic regulation but also be essential for developing targeted therapies that optimize healthspan and promote healthy aging in both sexes.

## Acknowledgements

This study was supported by Impetus, RF1AG078170, R01ES033171, CURES Center Grant P30ES020957, and CLEAR P42ES030991 for M.S. L.K was further supported by 5T32GM142519-02 and L.S was supported by 5T32HL120822-09. Services and support were provided by the WSU Microscopy, Imaging and Cytometry Resources Core P30CA22453 and R50CA251068-01 and WSU Genomics Core.

## Author contributions

H.S.M.J., L.K.D., L.S., A.T.D.S., L.K., R.S and S.S. carried out the research and reviewed the manuscript. M.S. designed the study and analyzed the data. M.S. wrote the manuscript and is responsible for the integrity of this work. All authors approved the final version of the manuscript.

## Declaration of interests

The authors declare no competing interests.

## Data Availability Statement

Further information and requests for resources and reagents should be directed to and will be fulfilled by the Lead Contact, Marianna Sadagurski. Any additional information required to reanalyze the data reported in this paper is available from the lead contact upon request.

## Notes

### Competing Interest Statement

The authors have declared no competing interest.

## REFERENCES

Bou Sleiman, M., Roy, S., Gao, A.W., Sadler, M.C., von Alvensleben, G.V.G., Li, H., Sen, S., Harrison, D.E., Nelson, J.F., Strong, R., Miller, R.A., Kutalik, Z., Williams, R.W. and Auwerx, J., 2022, Sex- and age-dependent genetics of longevity in a heterogeneous mouse population. Science, 377: eabo3191.

Coll, A.P. and Yeo, G.S., 2013, The hypothalamus and metabolism: integrating signals to control energy and glucose homeostasis. Curr Opin Pharmacol, 13: 970–976.

Dave, C.V., Schneeweiss, S., Kim, D., Fralick, M., Tong, A. and Patorno, E., 2019, Sodium-Glucose Cotransporter-2 Inhibitors and the Risk for Severe Urinary Tract Infections: A Population-Based Cohort Study. Ann Intern Med, 171: 248–256.

Evans, M.C., Rizwan, M.Z. and Anderson, G.M., 2014, Insulin action on GABA neurons is a critical regulator of energy balance but not fertility in mice. Endocrinology, 155: 4368–4379.

Grundy, S.M., 2004, Obesity, Metabolic Syndrome, and Cardiovascular Disease. The Journal of Clinical Endocrinology & Metabolism, 89: 2595–2600.

Han, E., Kim, A., Lee, S.J., Kim, J.Y., Kim, J.H., Lee, W.J. and Lee, B.W., 2018, Characteristics of Dapagliflozin Responders: A Longitudinal, Prospective, Nationwide Dapagliflozin Surveillance Study in Korea. Diabetes Ther, 9: 1689–1701.

Hierro-Bujalance, C., Infante-Garcia, C., Del Marco, A., Herrera, M., Carranza-Naval, M.J., Suarez, J., Alves-Martinez, P., Lubian-Lopez, S. and Garcia-Alloza, M., 2020, Empagliflozin reduces vascular damage and cognitive impairment in a mixed murine model of Alzheimer’s disease and type 2 diabetes. Alzheimers Res Ther, 12: 40.

Iorio, V., Festa, M., Rosati, A., Hahne, M., Tiberti, C., Capunzo, M., De Laurenzi, V. and Turco, M.C., 2015, BAG3 regulates formation of the SNARE complex and insulin secretion. Cell Death Dis, 6: e1684.

Jackson, A.U., Fornes, A., Galecki, A., Miller, R.A. and Burke, D.T., 1999, Multiple-trait quantitative trait loci analysis using a large mouse sibship. Genetics, 151: 785–795.

Jayarathne, H.S.M., Debarba, L.K., Jaboro, J.J., Ginsburg, B.C., Miller, R.A. and Sadagurski, M., 2022, Neuroprotective effects of Canagliflozin: Lessons from aged genetically diverse UM-HET3 mice. Aging Cell, 21: e13653.

Koepsell, H., 2020, Glucose transporters in brain in health and disease. Pflugers Arch, 472: 1299–1343.

Kullmann, S., Hummel, J., Wagner, R., Dannecker, C., Vosseler, A., Fritsche, L., Veit, R., Kantartzis, K., Machann, J., Birkenfeld, A.L., Stefan, N., Haring, H.U., Peter, A., Preissl, H., Fritsche, A. and Heni, M., 2021, Empagliflozin Improves Insulin Sensitivity of the Hypothalamus in Humans With Prediabetes: A Randomized, Double-Blind, Placebo-Controlled, Phase 2 Trial. Diabetes Care.

Lau, J. and Herzog, H., 2014, CART in the regulation of appetite and energy homeostasis. Front Neurosci, 8: 313.

Li, G., Zhang, Y., Wilsey, J.T. and Scarpace, P.J., 2005, Hypothalamic pro-opiomelanocortin gene delivery ameliorates obesity and glucose intolerance in aged rats. Diabetologia, 48: 2376–2385.

Lima J.B.M., D.L.K., Khan M., Ubah C., Didyuk O., Ayyar I., Koch I., Sadagurski M., 2020, Growth hormone receptor (GHR)-expressing neurons in the hypothalamic arcuate nucleus regulate glucose metabolism and energy homeostasis. BioRxiv, 10.1101/2020.08.17.254862.

Lin, B., Koibuchi, N., Hasegawa, Y., Sueta, D., Toyama, K., Uekawa, K., Ma, M., Nakagawa, T., Kusaka, H. and Kim-Mitsuyama, S., 2014, Glycemic control with empagliflozin, a novel selective SGLT2 inhibitor, ameliorates cardiovascular injury and cognitive dysfunction in obese and type 2 diabetic mice. Cardiovasc Diabetol, 13: 148.

Liu, T., Xu, Y., Yi, C.X., Tong, Q. and Cai, D., 2022, The hypothalamus for whole-body physiology: from metabolism to aging. Protein Cell, 13: 394–421.

Miller, R.A., Harrison, D.E., Allison, D.B., Bogue, M., Debarba, L., Diaz, V., Fernandez, E., Galecki, A., Garvey, W.T., Jayarathne, H., Kumar, N., Javors, M.A., Ladiges, W.C., Macchiarini, F., Nelson, J., Reifsnyder, P., Rosenthal, N.A., Sadagurski, M., Salmon, A.B., Smith, D.L., Jr., Snyder, J.M., Lombard, D.B. and Strong, R., 2020, Canagliflozin extends life span in genetically heterogeneous male but not female mice. JCI Insight, 5.

Nagai, K., Yoshida, S. and Konishi, H., 2014, Gender differences in the gene expression profiles of glucose transporter GLUT class I and SGLT in mouse tissues. Pharmazie, 69: 856–859.

Naznin, F., Sakoda, H., Okada, T., Tsubouchi, H., Waise, T.M.Z., Arakawa, K. and Nakazato, M., 2017, Canagliflozin, a sodium glucose cotransporter 2 inhibitor, attenuates obesity-induced inflammation in the nodose ganglion, hypothalamus, and skeletal muscle of mice. European Journal of Pharmacology, 794: 37–44.

Sawada, Y., Izumida, Y., Takeuchi, Y., Aita, Y., Wada, N., Li, E., Murayama, Y., Piao, X., Shikama, A., Masuda, Y., Nishi-Tatsumi, M., Kubota, M., Sekiya, M., Matsuzaka, T., Nakagawa, Y., Sugano, Y., Iwasaki, H., Kobayashi, K., Yatoh, S., Suzuki, H., Yagyu, H., Kawakami, Y., Kadowaki, T., Shimano, H. and Yahagi, N., 2017, Effect of sodium-glucose cotransporter 2 (SGLT2) inhibition on weight loss is partly mediated by liver-brain-adipose neurocircuitry. Biochem Biophys Res Commun, 493: 40–45.

Short, A.K., Thai, C.W., Chen, Y., Kamei, N., Pham, A.L., Birnie, M.T., Bolton, J.L., Mortazavi, A. and Baram, T.Z., 2023, Single-Cell Transcriptional Changes in Hypothalamic Corticotropin-Releasing Factor-Expressing Neurons After Early-Life Adversity Inform Enduring Alterations in Vulnerabilities to Stress. Biol Psychiatry Glob Open Sci, 3: 99–109.

Siklar, Z., Kontbay, T., Colclough, K., Patel, K.A. and Berberoglu, M., 2023, Expanding the Phenotype of TRMT10A Mutations: Case Report and a Review of the Existing Cases. J Clin Res Pediatr Endocrinol, 15: 90–96.

Snyder, J.M., Casey, K.M., Galecki, A., Harrison, D.E., Jayarathne, H., Kumar, N., Macchiarini, F., Rosenthal, N., Sadagurski, M., Salmon, A.B., Strong, R., Miller, R.A. and Ladiges, W., 2022, Canagliflozin retards age-related lesions in heart, kidney, liver, and adrenal gland in genetically heterogenous male mice. GeroScience.

Spanswick, D., Smith, M.A., Mirshamsi, S., Routh, V.H. and Ashford, M.L., 2000, Insulin activates ATP-sensitive K+ channels in hypothalamic neurons of lean, but not obese rats. Nat Neurosci, 3: 757–758.

Tahara, A., Takasu, T., Yokono, M., Imamura, M. and Kurosaki, E., 2016, Characterization and comparison of sodium-glucose cotransporter 2 inhibitors in pharmacokinetics, pharmacodynamics, and pharmacologic effects. J Pharmacol Sci, 130: 159–169.

van Gerwen, J., Shun-Shion, A.S. and Fazakerley, D.J., 2023, Insulin signalling and GLUT4 trafficking in insulin resistance. Biochemical Society Transactions, 51: 1057–1069.

van Ruiten, C.C., Veltman, D.J., Schrantee, A., van Bloemendaal, L., Barkhof, F., Kramer, M.H.H., Nieuwdorp, M. and RG, I.J., 2022, Effects of Dapagliflozin and Combination Therapy With Exenatide on Food-Cue Induced Brain Activation in Patients With Type 2 Diabetes. J Clin Endocrinol Metab, 107: e2590–e2599.

Williams, G., Bing, C., Cai, X.J., Harrold, J.A., King, P.J. and Liu, X.H., 2001, The hypothalamus and the control of energy homeostasis: different circuits, different purposes. Physiol Behav, 74: 683–701.

Wong, C.K.H., Tang, E.H.M., Man, K.K.C., Chan, E.W.Y., Wong, I.C.K. and Lam, C.L.K., 2021, SGLT2i as fourth-line therapy and risk of mortality, end-stage renal diseases and cardiovascular diseases in patients with type 2 diabetes mellitus. Diabetes & Metabolism, 47: 101196.

Xu, A.W., Kaelin, C.B., Takeda, K., Akira, S., Schwartz, M.W. and Barsh, G.S., 2005, PI3K integrates the action of insulin and leptin on hypothalamic neurons. J Clin Invest, 115: 951–958.

Xu, D., Chandler, O., Wee, C., Ho, C., Affandi, J.S., Yang, D., Liao, X., Chen, W., Li, Y., Reid, C. and Xiao, H., 2021, Sodium-Glucose Cotransporter-2 Inhibitor (SGLT2i) as a Primary Preventative Agent in the Healthy Individual: A Need of a Future Randomised Clinical Trial? Frontiers in Medicine, 8.

Yang, S.B., Tien, A.C., Boddupalli, G., Xu, A.W., Jan, Y.N. and Jan, L.Y., 2012, Rapamycin ameliorates age-dependent obesity associated with increased mTOR signaling in hypothalamic POMC neurons. Neuron, 75: 425–436.

Zhuang, A., Tan, Y., Liu, Y., Yang, C., Kiriazis, H., Grigolon, K., Walker, S., Bond, S.T., McMullen, J.R., Calkin, A.C. and Drew, B.G., 2023, Deletion of the muscle enriched lncRNA Oip5os1 induces atrial dysfunction in male mice with diabetes. Physiol Rep, 11: e15869.

