## Supplementary data for "Hypothalamic Sex-Specific Metabolic Shift by Canagliflozin during Aging"

**Running title:** Cana effect on hypothalamus in aging

**Keywords:** Canagliflozin, brain, hypothalamus, metabolism, longevity

***Corresponding author:**
**Marianna Sadagurski,**
**Department of Biological Sciences,**
**Integrative Biosciences Center**
**Wayne State University**
**Room 2418 IBio,**
**6135 Woodward, Detroit, MI 48202**


### Supplemental information

**Figure S1: Body composition and glucose metabolism in Cana-fed mice.** Body weight (A) and fat mass (B) after 4 weeks of Cana treatment. HOMA IR at 12 months of age for males (C) and females (D). Glucose tolerance test at 12 months of age in males (E) and females (F). Data represented as mean ± SEM, n= 4-6 mice/group. Two-way ANOVA followed by Newman-Keuls analysis (B-C, E-F, H-I, K-L, N-O, and Q-R). *p<0.05, **p<0.01, ***p<0.001. P values for the effect of diet, sex, and the interaction represent the significant p values from the two-way ANOVA.

**Figure S2: Energy homeostasis parameters measured in 12-month-old Cana-fed mice.** (A-F) Respiratory exchange ratio (RER) for (A-C) males and (D-F) females. (G-L) Locomotor activity in (G-I) males and (J-L) females. Food intake in (M) males and (N) females. Water intake in males (O) and females (P). Data represented as mean ± SEM, n= 5-6 mice/group. Two-way ANOVA followed by Newman-Keuls analysis (B-C, E-F, H-I, K-L, N-O, and Q-R). **p<0.01. P values for the effect of diet, sex, and the interaction during 24 hours represent the significant p values from the two-way ANOVA.

**Figure S3: Energy homeostasis parameters measured in 25-month-old Cana-fed mice.** (A-F) Oxygen consumption (VO2) in males (A-C) and females (D-F). (G-L) Carbon dioxide production (VCO2) in males (G-I) and females (J-L). (M-R) Locomotor activity in males (M-O) and females (P-R). Food intake in males (S) and females (T). Water intake in males (U) and females (V). Data represented as mean ± SEM, n= 5-6 mice/group. Two-way ANOVA followed by Newman-Keuls analysis (B-C, E-F, H-I, K-L, N-O, and Q-R). *p<0.05, **p<0.01. P values for the effect of diet, sex, and the interaction between diet and sex during the light and dark cycle represent the significant p values from the two-way ANOVA.

**Figure S4: Principal component analysis (PCA) of the hypothalamus isolated from Cana males and females.** Samples distribution of (A) 12-month-old male mice, (B) 12-month-old female mice, (C) 25-month-old male mice and (D) 25-month-old female mice. PCA was analyzed using the RStudio.Ink (version 2023.12.0369)

## S1

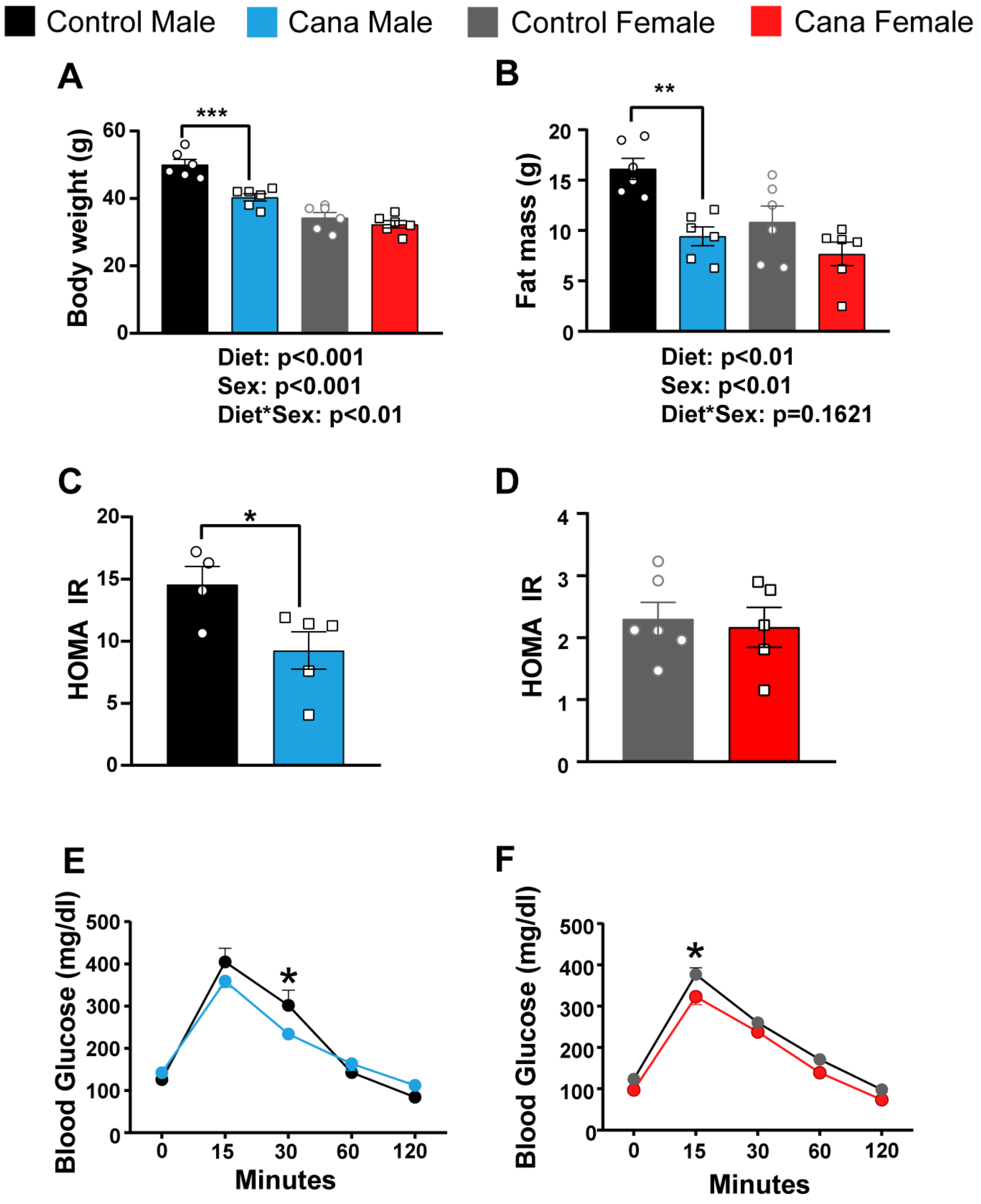

Control Male
Cana Male
Control Female
Cana Female
A
Body weight (g)
***
Diet: p<0.001
Sex: p<0.001
Diet*Sex: p<0.01
B
Fat mass (g)
**
Diet: p<0.01
Sex: p<0.01
Diet*Sex: p=0.1621
C
HOMA IR
*
D
HOMA IR
E
Blood Glucose (mg/dl)
*
Minutes
F
Blood Glucose (mg/dl)
*
Minutes

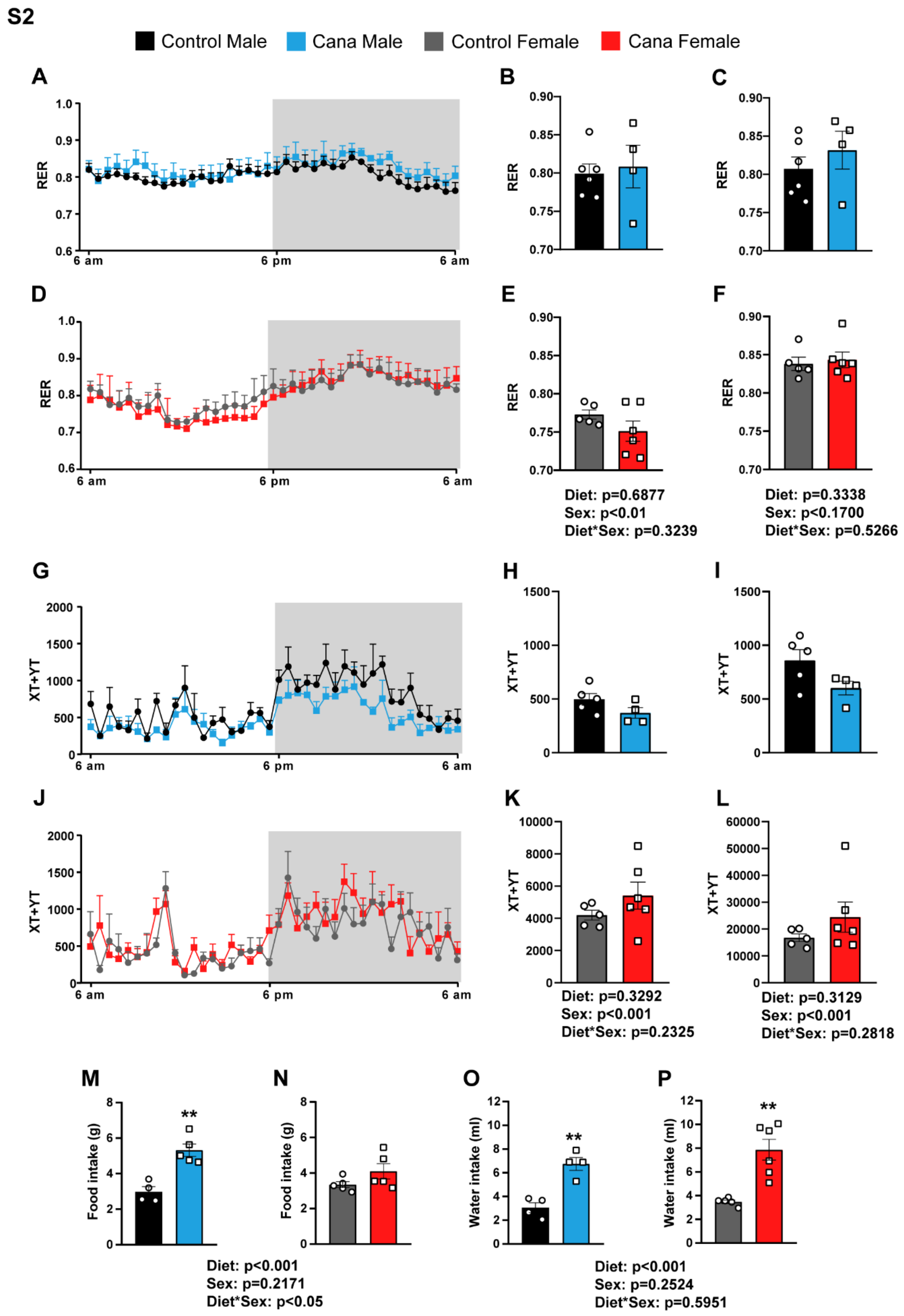
S2
Control Male
Cana Male
Control Female
Cana Female
A
B
C
D
E
F
RER
6 am
6 pm
6 am
Diet: p=0.6877
Sex: p<0.01
Diet*Sex: p=0.3239
Diet: p=0.3338
Sex: p<0.1700
Diet*Sex: p=0.5266
G
H
I
J
K
L
XT+YT
Diet: p=0.3292
Sex: p<0.001
Diet*Sex: p=0.2325
Diet: p=0.3129
Sex: p<0.001
Diet*Sex: p=0.2818
M
N
O
P
Food intake (g)
Water intake (ml)
**
Diet: p<0.001
Sex: p=0.2171
Diet*Sex: p<0.05
Diet: p<0.001
Sex: p=0.2524
Diet*Sex: p=0.5951

S3

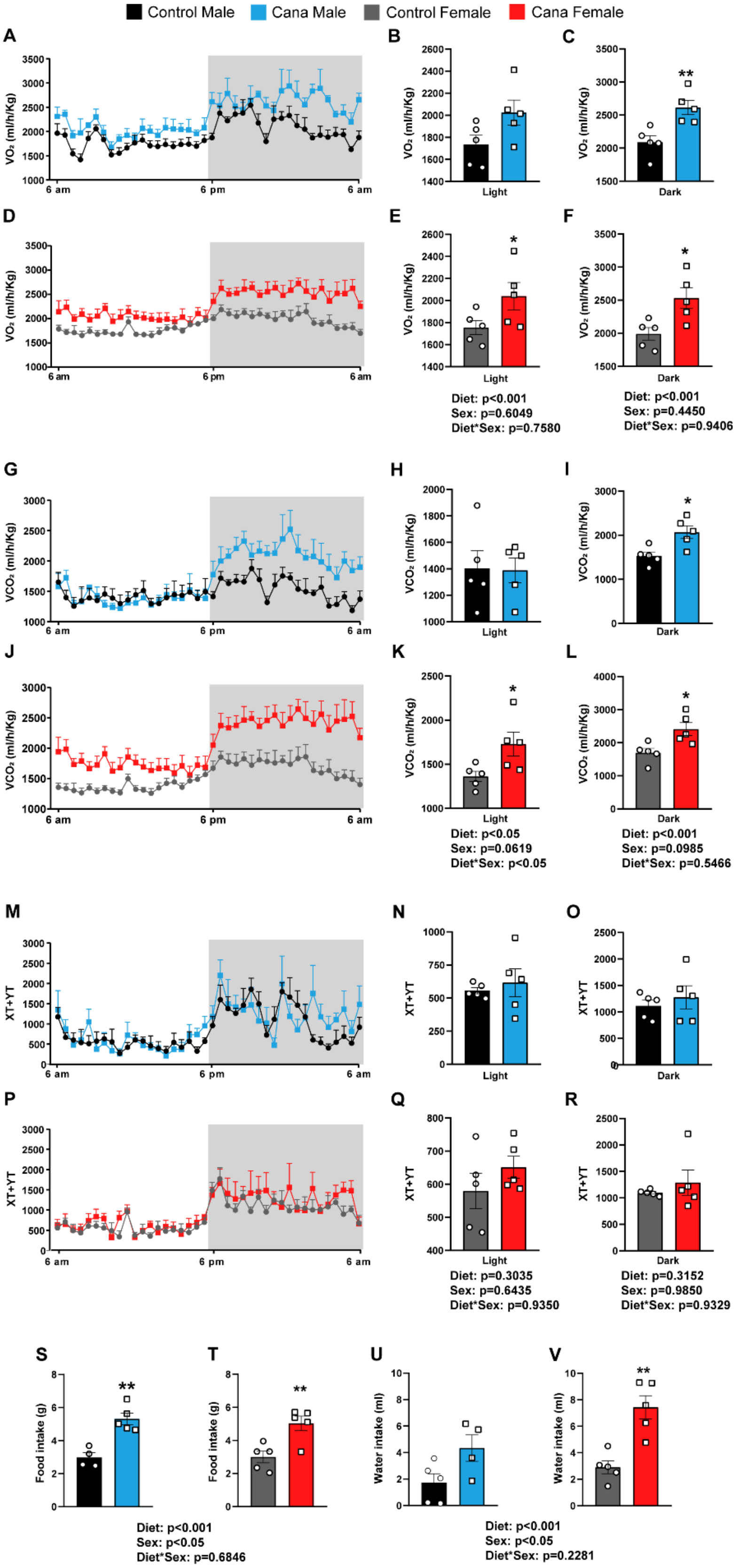

Control Male
Cana Male
Control Female
Cana Female
A
B
C
D
E
F
G
H
I
J
K
L
M
N
O
P
Q
R
S
T
U
V
VO₂ (ml/h/Kg)
VCO₂ (ml/h/Kg)
XT+YT
Food intake (g)
Water intake (ml)
6 am
6 pm
Light
Dark
Diet: p<0.001
Sex: p=0.6049
Diet*Sex: p=0.7580
Diet: p<0.001
Sex: p=0.4450
Diet*Sex: p=0.9406
Diet: p<0.05
Sex: p=0.0619
Diet*Sex: p<0.05
Diet: p<0.001
Sex: p=0.0985
Diet*Sex: p=0.5466
Diet: p=0.3035
Sex: p=0.6435
Diet*Sex: p=0.9350
Diet: p=0.3152
Sex: p=0.9850
Diet*Sex: p=0.9329
Diet: p<0.001
Sex: p<0.05
Diet*Sex: p=0.6846
Diet: p<0.001
Sex: p<0.05
Diet*Sex: p=0.2281

**S4**

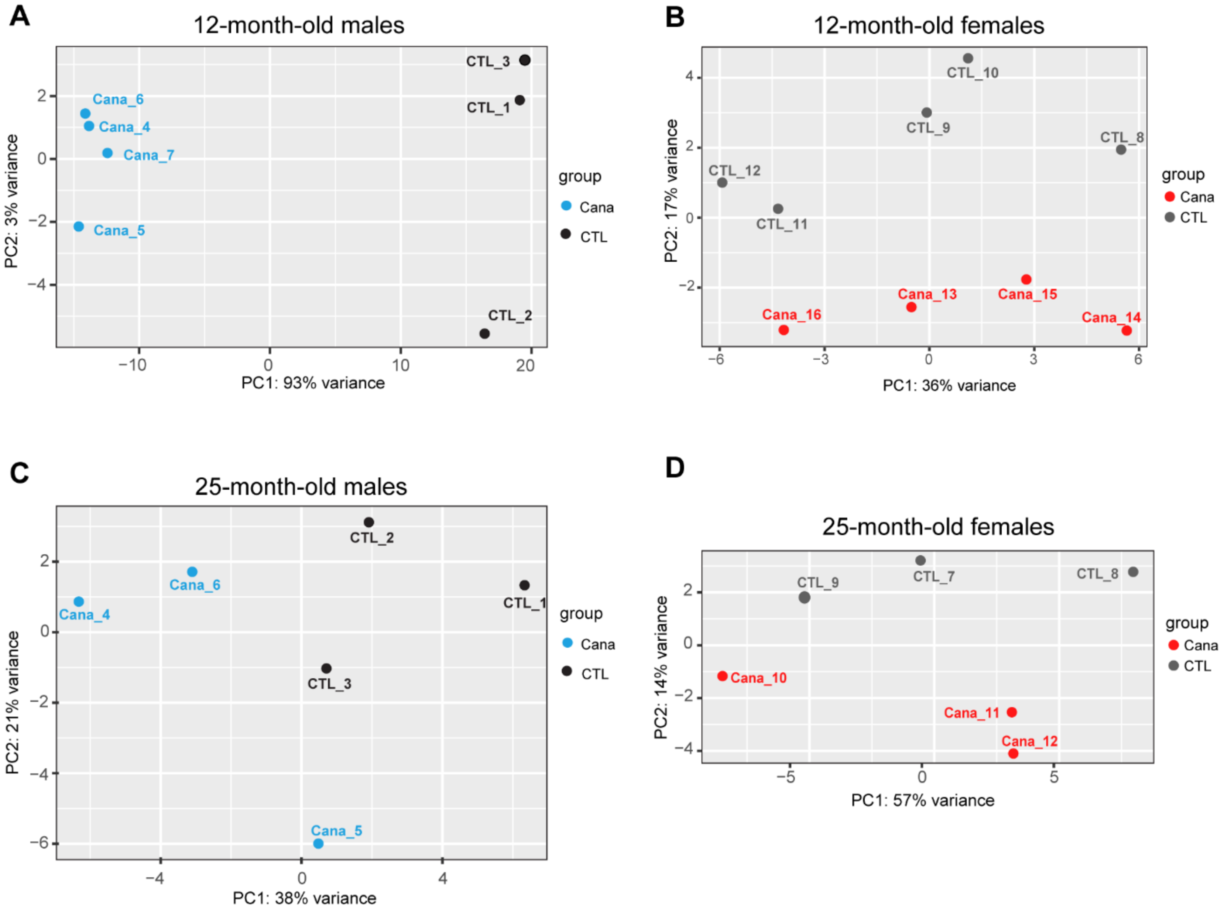
A
12-month-old males
PC1: 93% variance
PC2: 3% variance
Cana_6
Cana_4
Cana_7
Cana_5
CTL_3
CTL_1
CTL_2
group
Cana
CTL
B
12-month-old females
PC1: 36% variance
PC2: 17% variance
CTL_10
CTL_9
CTL_8
CTL_12
CTL_11
Cana_13
Cana_15
Cana_16
Cana_14
group
Cana
CTL
C
25-month-old males
PC1: 38% variance
PC2: 21% variance
CTL_2
Cana_6
CTL_1
Cana_4
CTL_3
Cana_5
group
Cana
CTL
D
25-month-old females
PC1: 57% variance
PC2: 14% variance
CTL_9
CTL_7
CTL_8
Cana_10
Cana_11
Cana_12
group
Cana
CTL
